# Complete Protection Against Lethal Crimean-Congo Hemorrhagic Fever Virus (CCHFV) Challenge in Mice Afforded by Immunization with Modified mRNA Encoding the Nucleocapsid Protein (NP) Alone

**DOI:** 10.1101/2025.04.21.649815

**Authors:** Sercan Keskin, Shaikh Terkis Islam Pavel, Rabia Sak, Fatemeh Bahadori, Ahmet Furkan Aslan, Münir Aktaş, Faruk Karakeçili, Ahmet Kalkan, Aykut Özdarendeli, Mehmet Ziya Doymaz

**Affiliations:** Department of Biotechnology, Institute of Health Sciences and Department of Microbiology, Beykoz Institute of Life Sciences and Biotechnology, Bezmialem Vakif University, Istanbul, Türkiye; Department of Microbiology, Faculty of Medicine and Vaccine Research, Development and Application Centre (ERAGEM), Erciyes University, Kayseri, Türkiye; Department of Pharmaceutical Biotechnology, Faculty of Pharmacy, Bezmialem Vakif University, Istanbul, Türkiye; Department of Analytical Chemistry, Faculty of Pharmacy, Cerrahpaşa University, Istanbul, Türkiye; Department of Parasitology, Faculty of Veterinary Medicine, University of Firat, Elazig, Türkiye; Department of Infectious Diseases and Clinical Microbiology, Faculty of Medicine, Erzincan Binali Yildirim University, Erzincan, Türkiye; Department of Infectious Diseases and Clinical Microbiology, Medical Faculty, Karadeniz Technical University, Trabzon, Türkiye; Department of Medical Microbiology, Medical School, and Beykoz Institute of Life Sciences and Biotechnology, Bezmialem Vakif University, Istanbul, Türkiye

## Abstract

**Background:** The Crimean-Congo Hemorrhagic Fever Virus (*Orthonairovirus haemorrhagiae*) causes a hemorrhagic fever with mortality rates reaching up to 40%. For years, this virus has maintained its position among the top priority pathogens identified by the World Health Organization (WHO). This is due to its endemic presence across a vast region—from Africa and Spain to the Balkans, the Middle East, and throughout Asia—its potential for human-to-human transmission, and the lack of an effective and approved vaccine or treatment. Therefore, the development of an effective vaccine against CCHFV is of critical importance. Building on the success of mRNA-based vaccines during the Coronavirus Disease 2019 (COVID-19) pandemic, this study reports the development of a messenger ribonucleic acid (mRNA) vaccine candidate expressing the nucleocapsid protein (NP) of CCHFV. The CCHFV NP in vitro transcript (IVT) was designed with pseudouridine (Ψ) nucleoside modification.

**Methods:** As part of the preclinical characterization of the IVT vaccine candidate, the biochemical and immunological properties of NP were confirmed in Huh-7 cells transfected with IVT NP-ΨmRNA. Afterwards, the efficacy of IVT NP-ΨmRNA immunization was evaluated in immunocompetent BALB/c and transiently immunosuppressed (IS) C57BL/6 mice. In CCHFV challenge studies, IS C57BL/6 mice were used. IS C57BL/6 mice were immunized intramuscularly with 2 doses of NP-ΨmRNA, either naked or encapsulated in Poly(lactic-co-glycolic acid) (PLGA) nanoparticles, administered 14 days apart.

**Findings:** High levels of CCHFV NP-specific humoral (IgM and IgG) and cellular (cytokine and lymphoproliferative) responses were demonstrated in BALB/c mice immunized with IVT NP-ΨmRNA. In challenge experiments, 100% protection was achieved by both the naked and PLGA-encapsulated IVT NP-ΨmRNA immunizations. Additionally, full protection was observed in mice immunized with inactivated CCHFV, whereas only 20% protection was detected in the unmodified IVT NP-mRNA vaccinated animals. In the protected mice, viral clearance was observed in the spleen, liver tissues, and blood on day 14 post-challenge.

**Interpretation:** This study demonstrates that NP, the most abundant protein of the virus, is capable of providing full protection as a standalone vaccine candidate. Furthermore, our report represents a crucial milestone in identifying a future vaccine candidate and paves the way for subsequent clinical studies.

**Fundings:** This study was supported by the Bezmialem Vakıf University Scientific Research Project BAP#20220402. Additionally, this study received support from TÜBİTAK 2211/C.

**Graphical Abstract:** 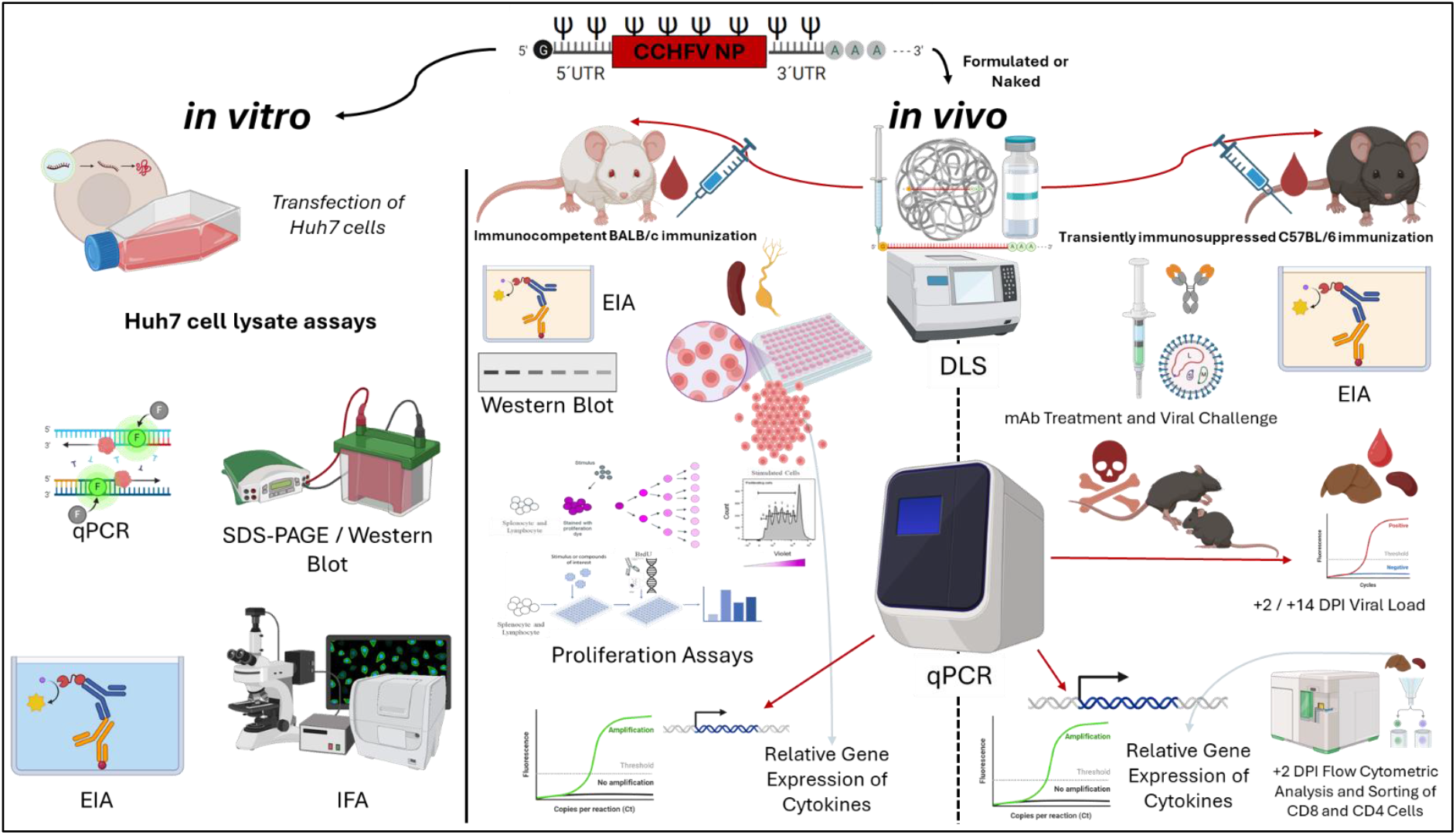

## Introduction

The Crimean-Congo hemorrhagic fever virus (CCHFV) is a significant pathogen transmitted to humans via arthropod vector ticks(1). Once a sporadic disease characterized by hemorrhagic fever, the early 2000s saw a marked increase in annual case reports, with mortality rates approaching 40%, and this brought the virus to the attention of global health authorities(2–4). Subsequent investigations showed that the vector population is endemic across a broad geographical range, extending from Japan to the Caucasus, to the Balkans and Southern Europe, including Spain, and throughout Africa(5–7). The widespread distribution of the virus, coupled with the potential expansion of vector habitats due to climate change, has positioned CCHFV as an emerging viral pathogen. Currently, the WHO has prioritized CCHFV as a critical pathogen requiring further research(8). No effective antiviral treatments nor approved vaccines exist to combat CCHFV infections(1, 9). Moreover, the virus’s association with resource-poor regions and the necessity for Biosafety Level 3 (or higher) containment for research have relegated CCHFV to the category of neglected pathogens.

In the present report, an mRNA based vaccine candidate against CCHFV is evaluated. Pre-COVID-19 era, mRNA-based vaccines were extensively tested, particularly against tumors and their development accelerated following the success of mRNA vaccines against SARS-CoV-2 during the pandemic(10). The widespread use of Moderna and Pfizer BioNTech’s mRNA vaccines (legal names; *Spikevax*-formerly referred to as “mRNA-1273”, and *Comirnaty* – formerly “BNT162b2”, respectively) and their role in rapidly mitigating the devastating impact of pandemic, have opened avenues for applying this technology to other viral pathogens, including CCHFV(11–14). Here, mRNA sequences encoding CCHFV NP, modified with pseudouridine, were utilized as immunizing antigen. These mRNAs were encapsulated in biodegradable Poly(lactic-co-glycolic acid) (PLGA) carrier particles and administered to here, mRNA sequences encoding CCHFV NP, modified with pseudouridine, were utilized as immunizing antigen. These mRNAs were encapsulated in biodegradable Poly(lactic-co-glycolic acid) (PLGA) carrier particles and administered to mice(15–17). The mice were experimentally immunosuppressed before challenge, and the effects of immunization on infection outcomes were assessed. The results demonstrate that the pseudo-uridine-modified IVT NP-ΨmRNA provided complete protection against viral challenge in the experimental model, indicating that a robust and effective vaccine candidate against CCHFV −0 a serious public health threat and neglected pathogen - is feasible. Our findings provide a comprehensive immunological analysis of IVT NP-ΨmRNA-PLGA and naked IVT NP-ΨmRNA vaccines in the context of CCHFV.

## Methods

### Biosafety and Ethical Declarations

All procedures involving infectious CCHFV were conducted under Animal Biosafety Level 3 (ABSL3) conditions, in accordance with the procedures approved by the Institutional Biosafety Committee of Erciyes University Laboratories in Kayseri. Animal experiments were approved by the Local Ethics Committee for Animal Experiments of Bezmialem Vakif University (protocol #2022/17, #2022/17-1 and #2022/89) and conducted under the supervision of a veterinarians and experienced personnel. The mice used in the challenge experiments were housed in HEPA-filtered cage systems in groups to acclimate them to ABSL3 conditions before the start of the challenge experiment. Nesting materials, food, and water were provided ad libitum. The surviving mice used in the experiments were humanely euthanized via cervical dislocation for the collection of necessary tissue and body fluids.

### Cells and Virus

Vero E6 (ATCC CRL-1586) cells were maintained in Dulbecco’s Modified Eagle Medium (DMEM) supplemented with 10% heat-inactivated fetal bovine serum (FBS), 100 mM L-glutamine, 50 U/mL penicillin, and 50 μg/mL streptomycin (Sigma-Aldrich, #P0781) at 37°C in a 5% CO_2_ environment(18). Human Hepatocellular Carcinoma cells (Huh-7) (JCRB- Japanese Collection of Research Bioresources Cell Bank-#0403) were grown in DMEM supplemented with 10% heat-inactivated FBS (Gibco, Brazil, #10270106), 100 U/mL penicillin, and 100 μg/mL streptomycin (Sigma-Aldrich, #P0781) at 37°C in a 5% CO_2_ environment(19). CCHFV Turkey-Kelkit06 strain was used the challenge experiments(18). The virus was propagated, titrated using pseudo-plaque assay (PPA) and stored as described earlier(18, 20). All viral procedures were carried out in a certified biosafety level 3 enhanced facility (BSL-3).

### In Vitro Transcription and Purification of ΨmRNA/mRNA

The S segment (GQ337053) of the CCHFV Turkey-Kelkit06 strain was placed downstream of the T7 promoter sequence of the designed pBILSAB WT plasmid (Figure S1) (appendix p 10). The S segment within the newly developed plasmid (pBILSAB WT/S) was unoptimized and contained dual 5’ and 3’ UTR sequences. After linearization with NdeI (NEB, #R0111L), the plasmid was used as a template. NP-ΨmRNA/mRNA was synthesized using the HiScribe® T7 ARCA mRNA Kit (with tailing) (NEB, #E2060S). Unrelated Luciferase ΨmRNAs were also synthesized as controls. During ΨmRNA synthesis, 10 mM β-pseudouridine (Cayman Chemical, #23383, Ann Arbor, MI, USA) was added and the final concentration was calculated as 1.25 mM. The synthesized mRNAs had Anti-Reverse CAP Analogs (ARCA) and were capped during synthesis by adding to the 5’ end. After synthesis, DNase I (RNase-free) provided in the kit was used to remove remaining DNA templates. The DNA-dependent RNA polymerization continued for 45 minutes at 37°C. This step was followed by the addition of an approximately 150-nucleotide Poly(A) tail, following the manufacturer’s instructions (NEB, UK). The reaction, completed with *E. coli* Poly(A) Polymerase provided in the kit, was purified by precipitating the mRNAs with lithium chloride (LiCl). The kit included LiCl solution for the rapid recovery of the synthesized mRNAs. LiCl precipitation effectively removed most unused NTPs and enzymes. LiCl was added to half of the reaction volume and incubated at −20°C for 30 minutes. RNA was pelleted by bench top centrifugation at 13000 g - for 15 minutes at 4°C. The supernatant was carefully discarded, and the pellet was washed with 500 μL of cold 70% ethanol and centrifuged again for 10 minutes at 4°C. Ethanol was carefully removed, and the tube was spun down to draw the remaining liquid to the bottom. The residual liquid was carefully removed using a sharp tip, and the pellet was air-dried and resuspended in ultrapure RNase-free water. To ensure complete dissolution of the RNA, it was heated at 65°C for 5-10 minutes. After thoroughly mixing, the RNA was stored at −80°C. The RNA concentration was determined by measuring the absorbance of ultraviolet light at 260 nm. Free nucleotides from the transcription reaction were removed by cleaning with LiCl and RNA concentrations were determined by spectrophotometer (Nanodrop™, Thermo Fisher Scientific, Pittsburgh, PA, USA).

A 1% agarose gel was used to analyze RNA samples. Agarose gel (Sigma-Aldrich) is prepared in 1X TAE (Tris-Acetate-EDTA) and poured into a comb mold. It is recommended to wash the equipment with RNase-free water at this stage. RiboRuler High Range RNA Ladder (Thermo Scientific, #SM1821) was used as the ladder. The ladder was mixed with 2X RNA Loading Dye containing Xylene Cyanol FF, Bromophenol Blue, and Ethidium Bromide. After heating the mixture at 70°C for 10 minutes, it was placed on ice for 3 minutes. The prepared agarose gel was loaded with the samples alongside the ladder. The gel tank was filled with 1X TAE. The ladder was similarly mixed with 2X RNA Loading Dye, heated, and placed on ice. The samples were run for 30 minutes at 40 volts, followed by 50 minutes at 80 volts. Images were captured with a UV imaging device (BioRad, Gel Doc XR+), and the analysis was performed using Image Lab software(21).

### Transfection of Huh-7 Cells and qPCR

In vitro transcribed and purified IVTs (ΨmRNA/mRNA) were transfected into Huh-7 cells using Lipofectamine 3000 (Invitrogen, #L3000008) and Xfect RNA Transfection Reagent (Takara, #631450). QPCR datas were normalized to GAPDH or β-actin using the 2–ΔΔCt method(22). Experiments were designed according to the manufacturer’s instructions as described in *Supplementary Method 1* of the appendix (p 3).

### Indirect Immunofluorescence Assay of mRNA Transfected cells

An IIFA (Indirect Immunofluorescence Assay) was performed to visualize the localization of the protein produced by Huh-7 cells transfected with mRNAs encoding the CCHFV NP protein, at 6, 24, and 48 hours post-transfection as described in *Supplementary Method 2* of the appendix (pp 3-4).

#### In-house EIA and Western Blot Assays

To assess the NP specific antibody responses qualitatively and quantitatively (antibody titers) in serum samples of immunized Balb/C and C57BL6 mice in-house developed EIA’s were utilized. For comparison, the samples were also tested using a commercial EIA (VectorBest, #D5052). The assay procedures were essentially similar to described earlier with modifications related to this study (23, 24). In house EIA’s were also utilized to assess the avidity of antibodies specific for NP. For this purpose, potassium thiocyanate (KSCN) displacement EIA method was applied as described earlier (25, 26) and also in *Supplementary Method 3* of the appendix (pp 4-6).

### PLGA Formulation of IVTs and DLS Measurements

In this study, PLGA nanoparticles (Sigma-Aldrich, #P2191) were used for formulation of antigens as described in *Supplementary Method 4* of the appendix (p 6)(16, 17, 27–29).

### BALB/c Mouse Immunizations

To evaluate the immunogenicity of the IVT structures, immunocompetent BALB/c mice were used. In this experiment, a total of 24 female BALB/c mice (8-12 weeks old) were randomly divided into six groups: (1) IVT NP-ΨmRNA-PLGA, (2) naked IVT NP-ΨmRNA, (3) IVT NP-mRNA-PLGA, (4) rNP + Alum, (5) iCCHFV + Alum, and (6) Sham (IVT Luc-ΨmRNA-PLGA + Alum). Each group was immunized a total of three times at 14-day intervals as described in *Supplementary Method 5* of the appendix (p 6).

### Collection and Culturing of BALB/c Mouse Spleen and Lymph Node Cells

The collection, culture, *in vitro* immune stimulation and labelling with bromodeoxyuridine (BrdU) of BALB/c spleen and lymph node cells were performed as described elsewhere(23, 24).

### Lymphoproliferation Assay (LPA)

Lymphoproliferation assay by BrdU labelling and by flow cytometric methods were performed to asses in vitro lymphocyte responses to the antigens as described earlier(23, 24) and also detailed *Supplementary Method 6* of the appendix (p 7). Following LPA, cytokine mRNAs produced by the stimulated lymphocytes were also determined by qPCR as described in *Supplementary Method 7* of the appendix (pp 7-8).

### C57BL/6 Mouse Immunizations and Challenge Experiments

C57BL/6 mouse model was used to determine the protective efficacy of IVT constructs and analyze the comparative protective response with other vaccine candidates. In this experiment, a total of 48 different-sex C57BL/6 mice (8-12 weeks old) were randomly divided into six groups: (1) IVT NP-ΨmRNA-PLGA, (2) naked IVT NP-ΨmRNA, (3) IVT NP-mRNA-PLGA, (4) rNP + Alum, (5) iCCHFV + Alum, and (6) Sham (IVT Luc-ΨmRNA-PLGA + Alum). Each group was immunized twice in total at 14-day intervals. The details of the immunizations are provided in *Supplementary Method 8* of the appendix (p 8). For challenge experiments and post-challenge evaluations, the procedures were as described earlier (30, 31).

### Determination of Viral Load

The viral loads in various tissues following challenge were determined by qPCR using primers 3’ Open Reading Frame (ORF) sequence of the S segment(23, 24). The procedures for determination of viral load in the tissues were as described in *Supplementary Method 9* of the appendix (pp 8-9).

### Determination of Cytokine mRNAs

Liver and spleen tissues from immunosuppressed (IS) C57BL/6 mice were collected at 2 dpi and 14 dpi time points and stored at −80°C for testing. Cytokine mRNA species purified from stored tissue samples were determined by qPCR essentially as described above and in detail in *Supplementary Method 10* of the appendix (p 9).

### Statistics

All graphs were generated using GraphPad Prism version 9.5.1 (GraphPad Software Inc., San Diego, CA, USA; www.graphpad.com). Data were considered statistically significant when p <0.05. Molecular biology procedures were simulated using SnapGene Viewer software (www.snapgene.com). Clinical scores, body weight, and body temperature were compared between groups. Survival analysis and the time to terminal disease after CCHF virus challenge were assessed using the log-rank (Kaplan-Meier) test. EIA results between groups were analyzed using two-way ANOVA with Bonferroni’s multiple comparison tests. Differences in viral levels were determined by a one-tailed Mann-Whitney test. Anti-nucleocapsid specific antibody avidity data and intracellular cytokine mRNA levels between groups were evaluated using two-way ANOVA (with Sidak’s correction). All data are presented as mean ± SD obtained from independent experiments. For the analysis of RT-PCR results, Ct values measured for each gene were normalized using the GAPDH gene with the Rotor-Gene® Q (Qiagen, USA). Relative expression levels were analyzed using the 2^-ΔΔCt method. EIA data were calculated by taking the average of absorbance measurements obtained from repeated samples. The cutoff value was calculated using the formula: average absorbance of each negative sample + 2 SD.

## Results

### Production of In Vitro Transcription (IVT) Products

The S segment of the CCHFV Turkey-Kelkit06 strain (GenBank No: GQ337053) was inserted into a plasmid to serve as a template for in vitro transcription (IVT). The plasmid map was generated using SnapGene software (Figure S1) (appendix p 10). The IVT NP-ΨmRNA and IVT NP-mRNA products produced via in vitro transcription were analyzed comparatively using non-denaturing agarose gel electrophoresis. Both products were expected to be around ~1.8 kb in size, but the addition of a Poly(A) tail increased the length by approximately 150 bases. The gel images showed that the IVT NP-ΨmRNA with base modifications and the IVT NP-mRNA produced with standard bases exhibited similar sizes and purity (Figure 1). The regions marked with red lines on the gel represent the expected IVT product sizes for both products.

**Figure 1.**
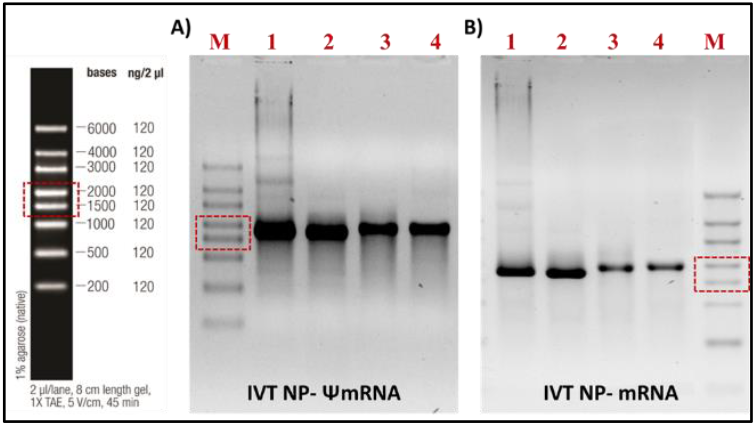
Production of modified and unmodified IVTs and electrophoresis on non-denaturing agarose gel. The expected size of the IVT products is approximately 1.8 kb. The addition of the poly(A) tail extends this by approximately 150 bases. **A)** IVT NP-ΨmRNA samples. **B)** IVT NP-mRNA samples. M: RiboRuler High Range RNA Ladder; 1: Samples taken immediately after the completion of the IVT reaction (1 μl from the reaction tube/well); 2: Samples taken after the addition of DNase I to the IVT reaction (1 μl); 3: Samples taken after the addition of poly(A) to the IVT reactions; 4: IVTs purified with LiCl. Red-boxed-dashed lines indicate the expected size of the IVT products.

### Temporal Analysis of IVT NP-ΨmRNA Expression in Huh-7 Cells

After transfection of Huh-7 cells with IVT NP-ΨmRNA, it was observed that the IVTs degraded over time within the cells. Transfection was performed using Xfect RNA Transfection Reagent, and naked IVT NP-ΨmRNAs were also added to the cells under the same conditions for comparison. The transfection process involved adding 2 μg/ml of IVT to cells that had reached 80% confluency. Total RNA was isolated from the cells at 6, 24, and 48 hours, and qPCR was performed using S segment primers (Figure S2) (appendix p 11). After transfection of Huh-7 cells with IVT NP-ΨmRNA, increasing protein expression in the cytoplasm was observed over time. The cells were incubated for 48 hours post-transfection, and the results are presented in Figure 2. Accordingly, NP expression increased over time in the transfected cells (Figure 2A). Western blot analysisof transfected Huh-7 cells after 48 hours of transfection showed the desired band profiles only from cell lysates transfected with mRNA (Figure 2B). Indirect immunofluorescence assay (IIFA) was used to analyze NP expression in cells at 0, 6, 24, and 48 hours post-transfection. Again, overtime increasing expression of NP in the cells was observed (Figure 2C).

**Figure 2.**
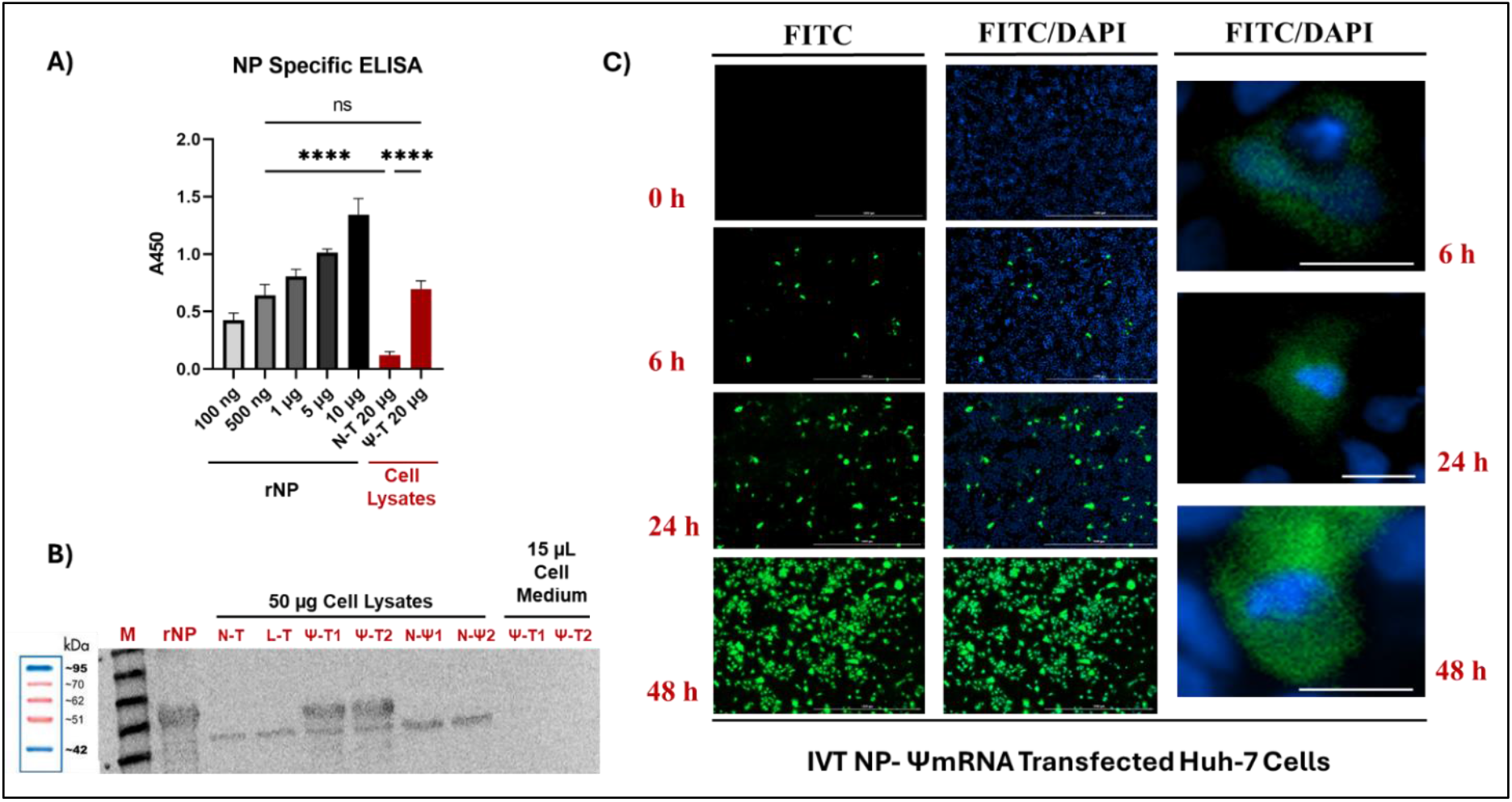
IVT NP-ΨmRNA is expressed in the cytoplasm of Huh-7 cells, showing an increase over time. **A)** Detection of NP in cell lysated transfected with IVT NP-ΨmRNA by EIA. Different amounts of rNP were coated on EIA plates. Forty-eight hours post-transfection, cells were lysed. Lysates from transfected Huh-7 cells (IVT NP-ΨmRNA (Ψ-T)) and non-transfected (N-T) Huh-7 cells were normalized using the Bradford Protein Assay and then coated onto the plates. The primary antibody used was serum from mice immunized with iCCHFV. **B)** Western blot analysis of expressed NP in cell lysated transfected with IVT NP-ΨmRNA. The primary antibody was serum from mice immunized with iCCHFV. L-T refers to transfection with IVT Luc-ΨmRNA, Ψ-T1 refers to transfection with Lipofectamine 3000, and Ψ-T2 refers to transfection with Xfect RNA Transfection Reagent. N-Ψ1 (2 μg/ml) and N-Ψ2 (6 μg/ml) refer to cells incubated with naked IVT NP-ΨmRNA added to the supernatant. **C)** IIFA of Huh-7 cells transfected with IVT NP-ΨmRNA. The cells were tested at 6, 24, and 48 hours post-transfection (0h) for expression localization and intensity using IIFA. The primary antibody was serum from mice immunized with iCCHFV, and the secondary antibody was FITC-conjugated Anti-Mouse IgG H&L. Increased NP expression was noticed over the 48-hour period. ns; P>0.05, ****P ≤ 0.0001 (One-way ANOVA).

### Immunization of BALB/c Mice with IVT NP-ΨmRNA Induces Strong NP-Specific Antibody Responses

The characterization of IVTs formulated with PLGA is performed. According to dynamic light scattering (DLS) results, IVTs formulated with PLGA had a negative zeta potential and formed particles of approximately 100 nanometers in size (Figure S3) (appendix p 12). Antibody responses to PLGA-formulated IVTs in BALB/c mice were tested following immunizations. Blood samples from mice were obtained after immunizations, and EIA results demonstrated that IVT NP-ΨmRNA-PLGA elicited a strong humoral response (Figure 3). NP-specific IgM response was highest in the blood taken 14 days after the first immunization specifically IVT NP-ΨmRNA group showing the highest readings (Figure 3A) On the other hand, the NP-specific IgG response increased with each immunization in blood samples taken at 14-day intervals. The IgG response was lowest for IVT NP-mRNA immunizations and highest for IVT NP-ΨmRNAs immunizations (Figure 3B). The accuracy of in-house tests was confirmed using a commercial test (13)(Figure 3C). The highest antibody titers in commercial assay were noticed in iCCHFV-immunized mice following the final immunization with. The avidity of NP-specific IgG antibodies increased with successive immunizations (Figure 3D), with the highest avidity levels observed in the IVT NP-ΨmRNAs group. The lowest avidity levels were observed in the IVT NP-mRNA immunized animals. Western blot analysis of sera from BALB/c mice immunized with IVT NP-ΨmRNA-PLGA after the third immunization showed that the antibodies reacted with both rNP and IVT NP-ΨmRNA-transfected cell lysates (Figure 3E). Serially diluted serum samples from immunized BALB/c mice were also tested and endpoint titers of these sera were determined (Figure S4) (appendix p 12). These results demonstrated that NP-specific antibodies could be detected even at 1:65536 dilutions of sera from IVT NP-ΨmRNAs immunized groups. Overall, the results of antibody assays indicated that immunization with IVT NP-ΨmRNAs in naked forms or in PLGA induced high-avidity antibody responses in BALB/c mice, and these antibodies were able to recognise to viral NP produced in *E. coli* or in IVT NP-ΨmRNA-transfected cell lysates.

**Figure 3.**
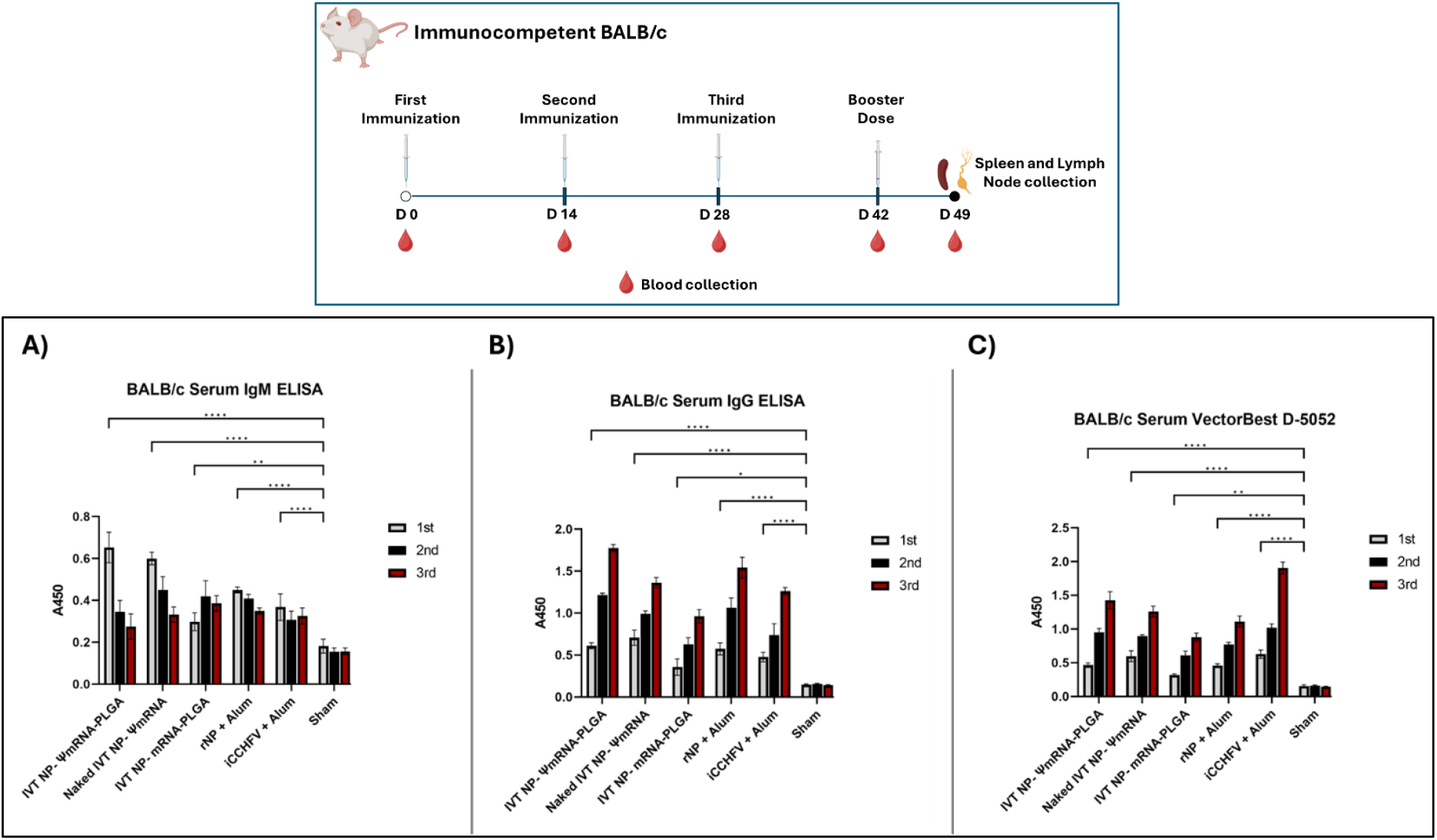
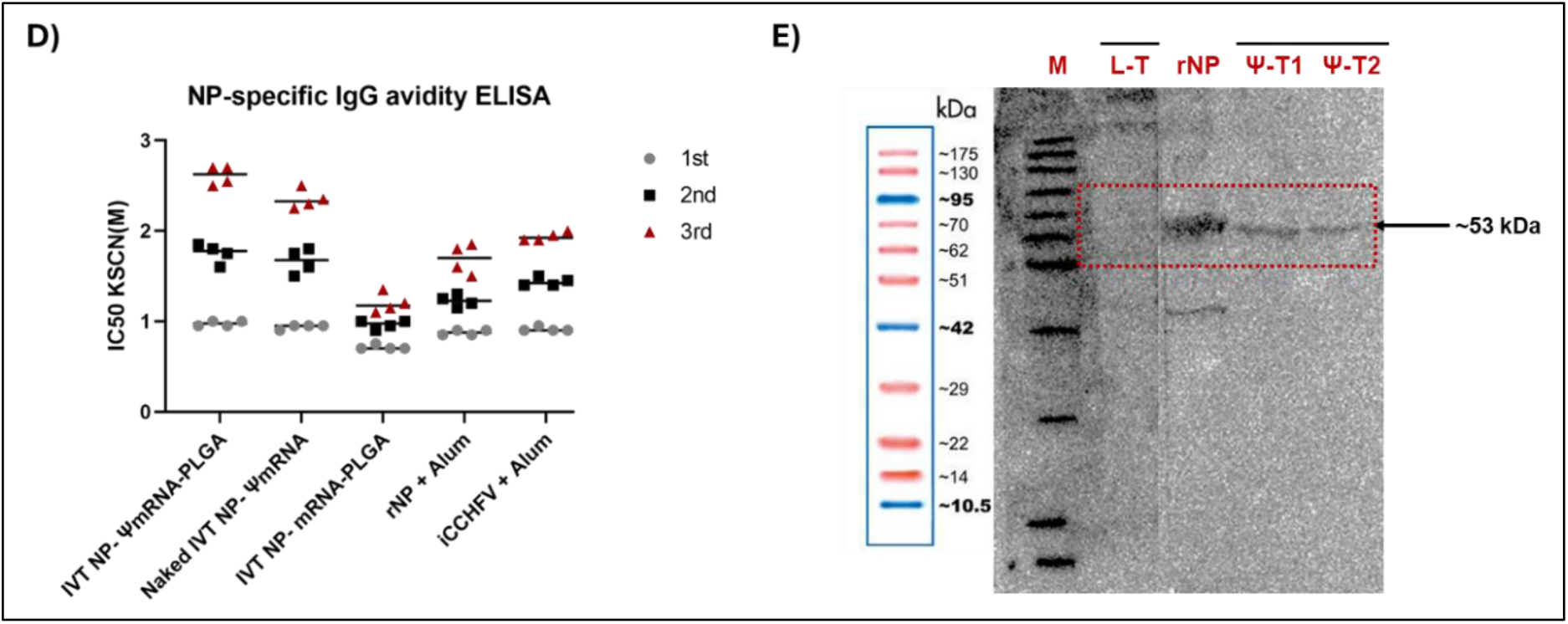
BALB/c mice immunized with IVT NP-ΨmRNA develop a strong humoral response and authentic antibodies. **A-B)** The EIA plate was coated with 10 μg rNP. **C)** The commercial assay (VectorBest D-5052) was used with modification of the secondary antibody for the model. **D)** The avidity of CCHFV NP-specific IgG responses was measured using a KSCN chemical displacement EIA. **E)** Forty-eight hours post-transfection, a portion of Huh-7 cells were lysed for Western blot analysis. Twenty-five μg of Huh-7 cell lysates (black-striped samples) were boiled in 6X Laemmli Buffer and loaded into SDS-PAGE wells. After blotting, the primary antibody used was serum from mice immunized three times with IVT NP-ΨmRNA-PLGA (1:100 dilution). Western blot results were visualized using ECL. L-T refers to transfection with IVT Luc-ΨmRNA, Ψ-T1 refers to transfection with Lipofectamine 3000, and Ψ-T2 refers to transfection with Xfect RNA Transfection Reagent.

### NP Stimulates Lymphocyte Proliferation In Vitro

To evaluate the cellular immune response, spleen and lymph node cells obtained from immunized mice were stimulated with NP in an in vitro environment, and the proliferative response of the cells was determined by flow cytometric analysis. In these tests, the positive control using ConA induced the highest cell division, while the specific stimulation with rNP generated a significant proliferation response in immune lymphocytes. Additionally, specific responses against iCCHFV and transfected cell lysates were also observed (Figure 4A-D). The index values of these responses were calculated, confirming the presence of a strong and specific T cell response (Figure 4E).

**Figure 4.**
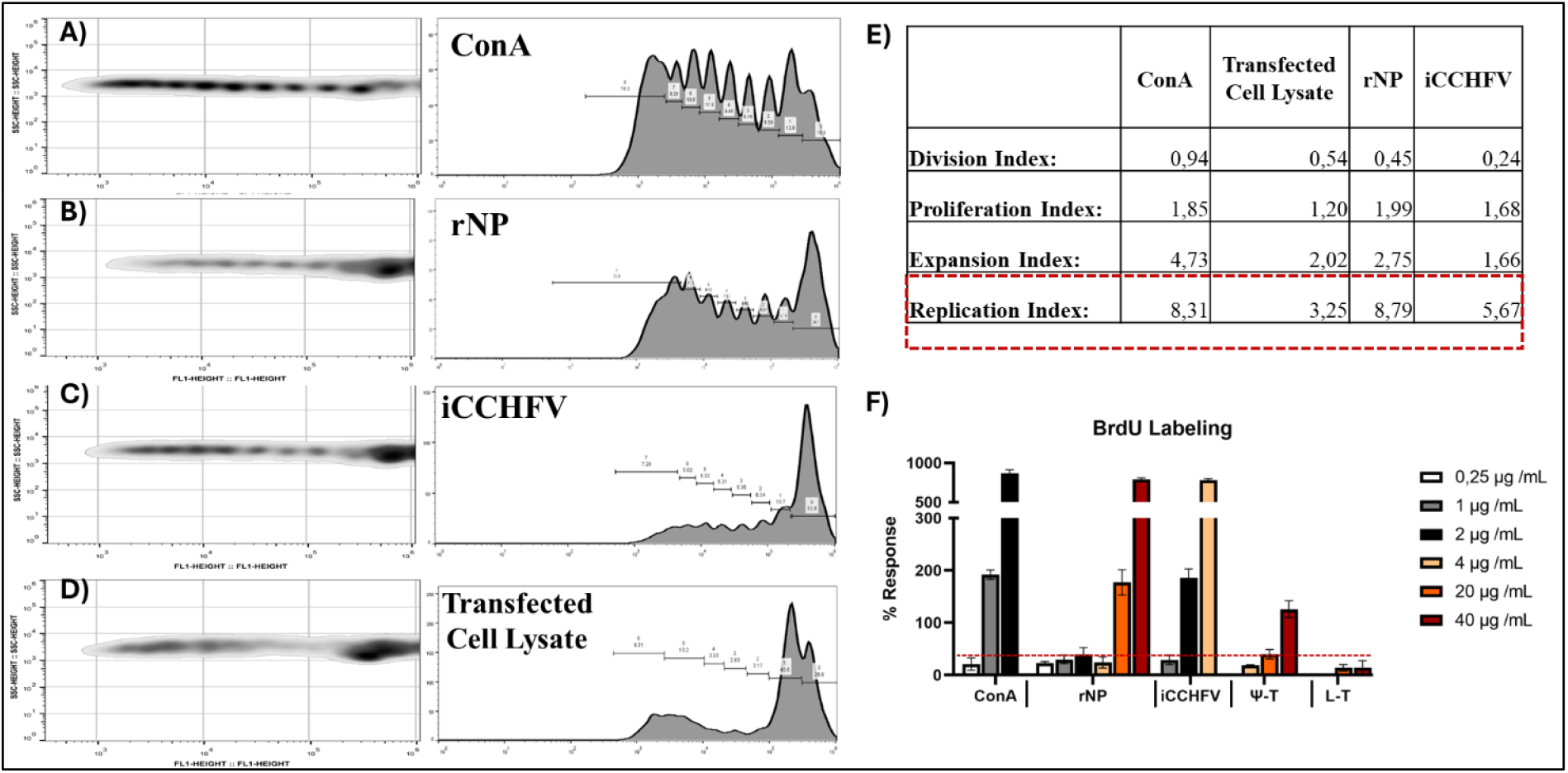
NP Stimulates Lymphocyte Proliferation In Vitro (Flow Cytometry Analysis). Spleens and lymph nodes (cervical, brachial, axillary, inguinal, lumbar, sacral, sciatic) were harvested from BALB/c mice immunized three times with IVT NP-ΨmRNA-PLGA and dissociated into single-cell suspensions. **A-D**. Co-cultures of 105 splenocytes and 105 lymphocytes were stimulated with ConA (positive control), rNP, iCCHFV, or IVT NP-ΨmRNA transfected cell lysates. Cells were labeled with CellTrace™ Violet before culturing, and flow cytometry analysis was performed 7 days post-stimulation. The first column shows Dot Blot data, and the second column displays the corresponding histograms, with each row representing the same set of data. **E**. Proliferation Response Index calculations were based on FlowJo data: **Division Index**: Total number of divisions / number of cells at the start of culture. **Proliferation Index**: Total number of divisions / cells that underwent division. **Expansion Index**: Total number of cells / cells at the start of culture. **Replication Index**: Total number of divided cells / cells that underwent division. **F**. Cells (2.0 × 105/well) were stimulated with varying concentrations of NP-containing components or ConA. After three days of incubation, cells were labeled with 10 μM BrdU for 24 hours, and the BrdU incorporation was detected colorimetrically at 490 nm. The percent response was calculated as follows: [(mean OD of antigen-stimulated group – mean OD of negative control group) / mean OD of negative control group] x 100. L-T refers to the lysates of cells transfected with IVT Luc-ΨmRNA, while Ψ-T denotes the lysates of cells transfected with IVT NP-ΨmRNA using Lipofectamine 3000 and Xfect RNA Transfection Reagent. Statistical significance was determined by two-way ANOVA (P ≤ 0.0001).

In addition to flow cytometry, lymphoproliferation experiments were conducted using BrdU labeling. This analysis was repeated with stimulatory components at different concentrations and found that NP-specific T cell responses showed higher rates of proliferation with increasing stimulatory concentrations (Figure 4F). It was also observed that the proliferating cells formed dense cell clusters (Figure S5) (appendix p 13). Following in vitro stimulation, the cytokines produced by the cells were also determined at the mRNA level. For this purpose, cytokine mRNA levels were compared to control cells containing RPMI-PBS, and relative gene expression values were calculated. These results particularly showed increased responses for IFN gamma, IL-12, and TNF-alpha (Figure S6) (appendix p 14). Significant increases were also detected in IL-2 and IL-15 cytokine responses.

### In Vivo Challenge Tests and Immune Correlates

Initial immunization and immune analyses conducted with Balb/c mice demonstrated that IVT NP-ΨmRNAs could elicit a successful immune responses. At this point, the studies were transferred to the C57BL/6 mouse model, where viral challenges could be conducted. First, C57BL/6 mice were immunized twice, and anti-NP antibodies were searched in serum samples. It was determined that all mice were immunized prior to challenge and reached high antibody titers (Figure S7) (appendix p 15).

In the C57BL/6 mouse model, the efficacy of immunization was tested in challenge experiments conducted under immunosuppressive conditions. For this purpose, immunized mice were intraperitoneally challenged with 400 ppfu Kelkit CCHFV and monitored for two weeks post-infection. The results showed that IVT NP-ΨmRNA-PLGA (n=6), Naked IVT NP-ΨmRNA (n=6), and iCCHFV (n=5) with two doses of immunization provided complete protection against lethal CCHFV challenge (Figure 5A). The animals in these groups had fully recovered by day 14. In contrast, animals immunized with unrelated mRNA vaccines succumbed to fatal infection by day 5. In the IVT NP-mRNA-PLGA group, only 20% (n=1) of the mice and in the rNP immunized group 40% (n=2) recovered by day 14.

**Figure 5.**
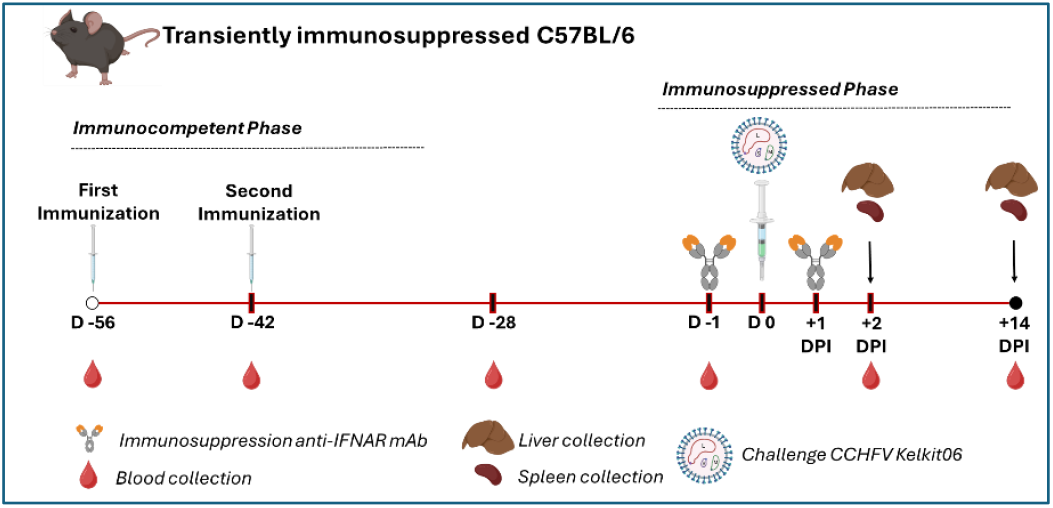
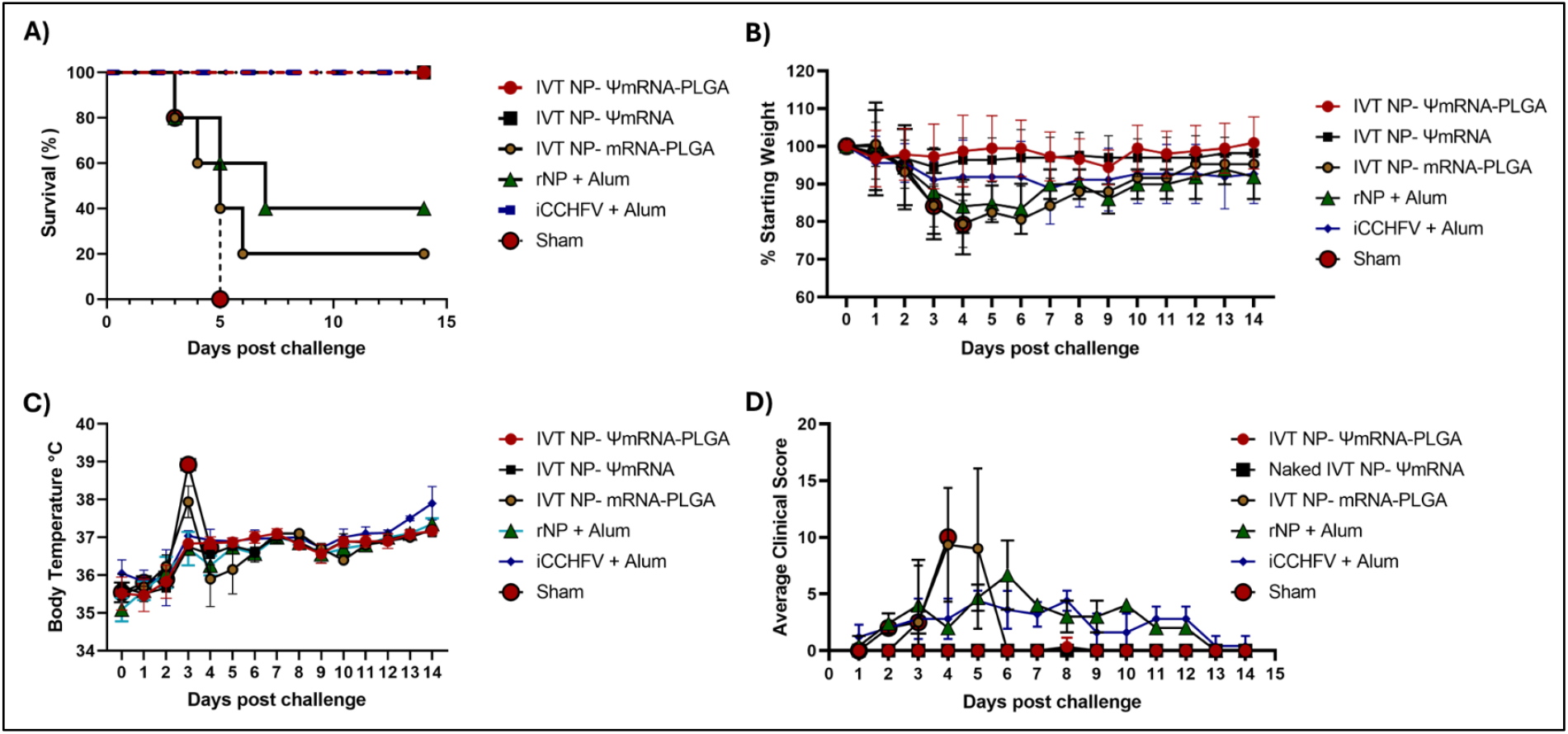
IVT NP-ΨmRNAs Provide Full Protection Against Lethal CCHFV Challenge in the C57BL/6 Mouse Model. C57BL/6 mice (n=48) were divided into six groups and immunized twice at 14-day intervals with IVT constructs, rNP, or iCCHFV, followed by a challenge with 400 PPFU of CCHFV Turkey-Kelkit06 strain under immunosuppressive conditions. Mice immunized with IVT NP-ΨmRNA-PLGA, naked IVT NP-ΨmRNA, and iCCHFV exhibited complete protection, with no severe clinical symptoms observed by day 14 post-infection **(A)**. Body weight **(B)**, temperature **(C)**, and clinical scores **(D)** were monitored.

The body weights (Figure 5B), body temperatures (Figure 5C), and clinical scores of the mice were monitored over the 14 days. These clinical scores are presented as the average for each group in Figure 5D. Additionally, the clinical scores of the mice in each group taken on different days were shown in a heat map format (Figure S8) (appendix p 16). Furthermore, in mice immunized with IVT NP-ΨmRNAs, in addition to full protection, no severe clinical symptoms were observed. It was noted that while inactive CCHFV provided 100% protection, it did not prevent the emergence of clinical symptoms.

### Viral Load and qPCR Analyses

During the challenge, blood, liver, and spleen samples from randomly selected mice from each group were collected on day 2 (+2 DPI), the period when viral infection was at its peak. Total RNA was extracted from these samples, and cDNA templates were analyzed in real-time quantitative PCR using CCHFV Kelkit strain NP-specific qPCR primers. In analyses performed with real-time PCR, viral loads (viral copy number/ml) were calculated based on the obtained Ct values. According to these results, high viral loads were detected in the blood, liver, and spleen tissues of negative controls and IVT NP-mRNA-PLGA immunized animals. In the rNP+Alum immunized group, although the viral load in the blood was low, high viral loads were detected in liver and spleen samples. By the end of the experiment (+14 DPI), complete viral clearance was observed in all the surviving mice (Figure 6).

**Figure 6.**
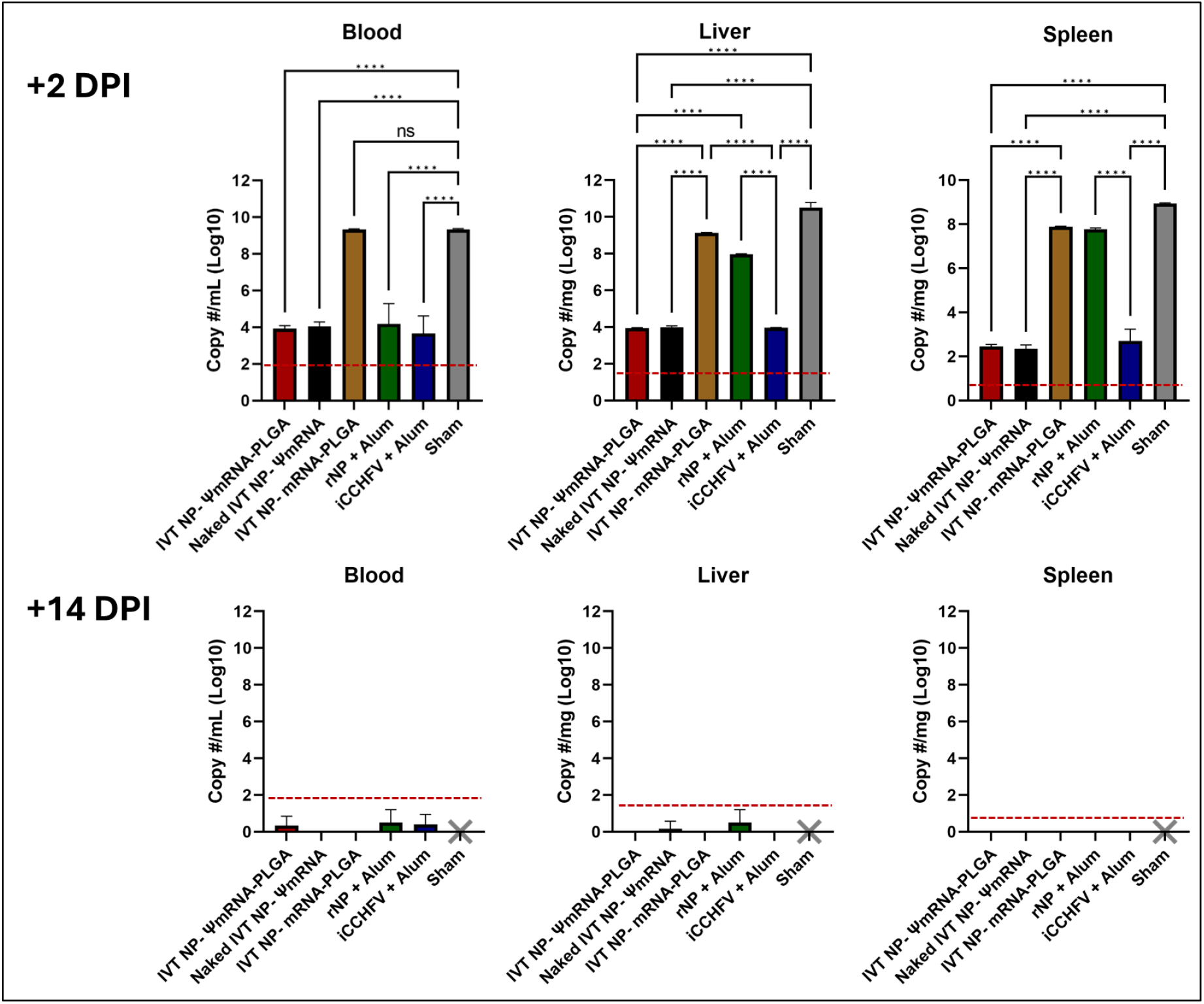
IVT NP-ΨmRNAs Lead to Complete Viral Clearance by 14 DPI in CCHFV Challenge. Total RNA was extracted from blood, liver, and spleen samples of C57BL/6 mice on day 2 (+2 DPI) and day 14 (+14 DPI) post-infection. Real-time quantitative PCR analysis was conducted using NP-specific qPCR primers for the CCHFV Kelkit strain. High viral loads were observed in the blood, liver, and spleen of negative control and IVT NP-mRNA-PLGA immunized mice. The rNP+Alum group exhibited low viral loads in the blood but high loads in the liver and spleen. Complete viral clearance was observed in all surviving mice by +14 DPI.

### Endpoint Titer and Avidity Analyses

In immunized mice, another immune parameter tested was the humoral immune response. For this purpose, NP-specific IgM and IgG responses of the mice were analyzed before (−1 DPI), during (+2 DPI), and after the challenge (+14 DPI) (Figure S9) (appendix p 17). Accordingly, high IgG antibody responses were noticed in animals immunized with IVT NP-ΨmRNA-PLGA and Naked IVT NP-ΨmRNA before and after the challenge (Figure S9A). In the experimental groups that showed full protection, OD data tended to be higher, whereas in groups where protection was less than complete, OD values tended to be lower. A similar trend was noted in IgM assays (Figure S9C). Additionally, high antibody levels were detected in surviving mice after the challenge. These results were confirmed with NP-based in-house and commercial IgG tests (Figure S9B).

In our study, serum antibody endpoint titers were also determined. Here, pooled animal serum was tested. The endpoint titers of IgG antibodies formed during the periods of −28 DPI, −1 DPI, +2 DPI, and +14 DPI after the second vaccination (−28 DPI) are presented in Figure 7A. Accordingly, high antibody titers were found in the groups providing full protection on days −28, −1, and +2 DPI. In all mice that survived after the challenge (+14 DPI), the antibody endpoint titers showed similarly high levels. Particularly, the mouse groups with low endpoint titers during the −28 DPI and −1 DPI periods were IVT NP-mRNA-PLGA, rNP+Alum and iCCHFV+Alum groups. During the challenge, only the iCCHFV group among these groups managed to survive with clinical symptoms. Endpoint titers during the challenge were also found to be low in mouse groups with low survival rates. Although the endpoint titers were somewhat lower in iCCHFV group mice, in the survived animals following the challenge, the titers increased.

**Figure 7.**
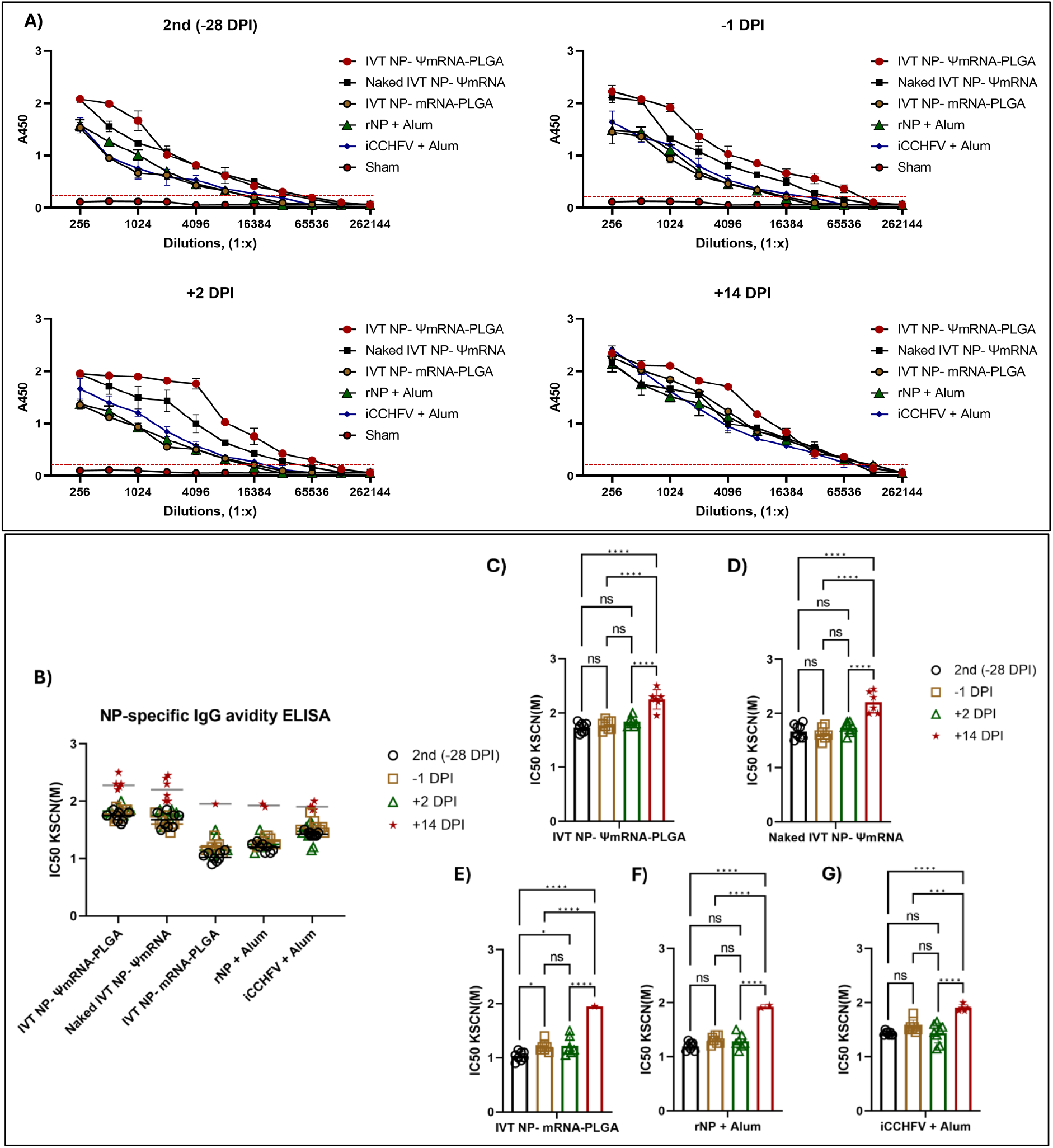
IVT NP-ΨmRNAs Induce High-Avidity Antibody Response with Elevated Endpoint Titers Post-Challenge. The avidity of NP-specific IgG antibodies was determined using the KSCN displacement assay. A) Endpoint titers were measured at multiple time points (−28 DPI, −1 DPI, +2 DPI, +14 DPI) following the second vaccination by EIA. High antibody titers were detected in IVT NP-ΨmRNA-PLGA and Naked IVT NP-ΨmRNA groups, correlating with full protection against CCHFV challenge. B) Antibody avidity was highest in the IVT NP-ΨmRNA-PLGA and Naked IVT NP-ΨmRNA groups before and after the challenge, while low-avidity groups improved avidity post-challenge. Figure 7C-G presents the detailed changes in antibody avidity, showing that even groups with initially low avidity successfully raised their response after the challenge.

This study also determined the avidity of antibodies stimulated in immunized mice (Figure 7B-G). Before the challenge, the antibodies with the highest avidity were found in the IVT NP-ΨmRNA-PLGA and Naked IVT NP-ΨmRNA mouse groups, while the antibody avidity results in other groups that did not show full protection remained low (Figure 7B). After the challenge, it was observed that all surviving mice produced antibodies with high avidity compared to their own groups (Figure 7B). The avidity levels indicated that the antibodies showed very high avidity, and these high values were again highest in the IVT NP-ΨmRNA-PLGA and Naked IVT NP-ΨmRNA mouse groups. Additionally, changes in antibody avidity were observed in the same group of mice before and during the challenge (Figure 7C-G).

### Analysis of Cellular Immune Response: Increase of CD3+ CD8+ Cells in Liver and Spleen

Alongside viral load tests conducted two days after the challenge, lymphocyte profiling was also performed on spleen and liver tissues, as shown in Figure 8. The relative proportions of CD4+ and CD8+ cells, compared to those in the sham group, were determined through flow cytometric measurements. Accordingly, a notable increase of up to 30% in CD8+ cells was observed, particularly in mouse groups with a 100% survival rate (Figure S10) (appendix p 18).

**Figure 8.**
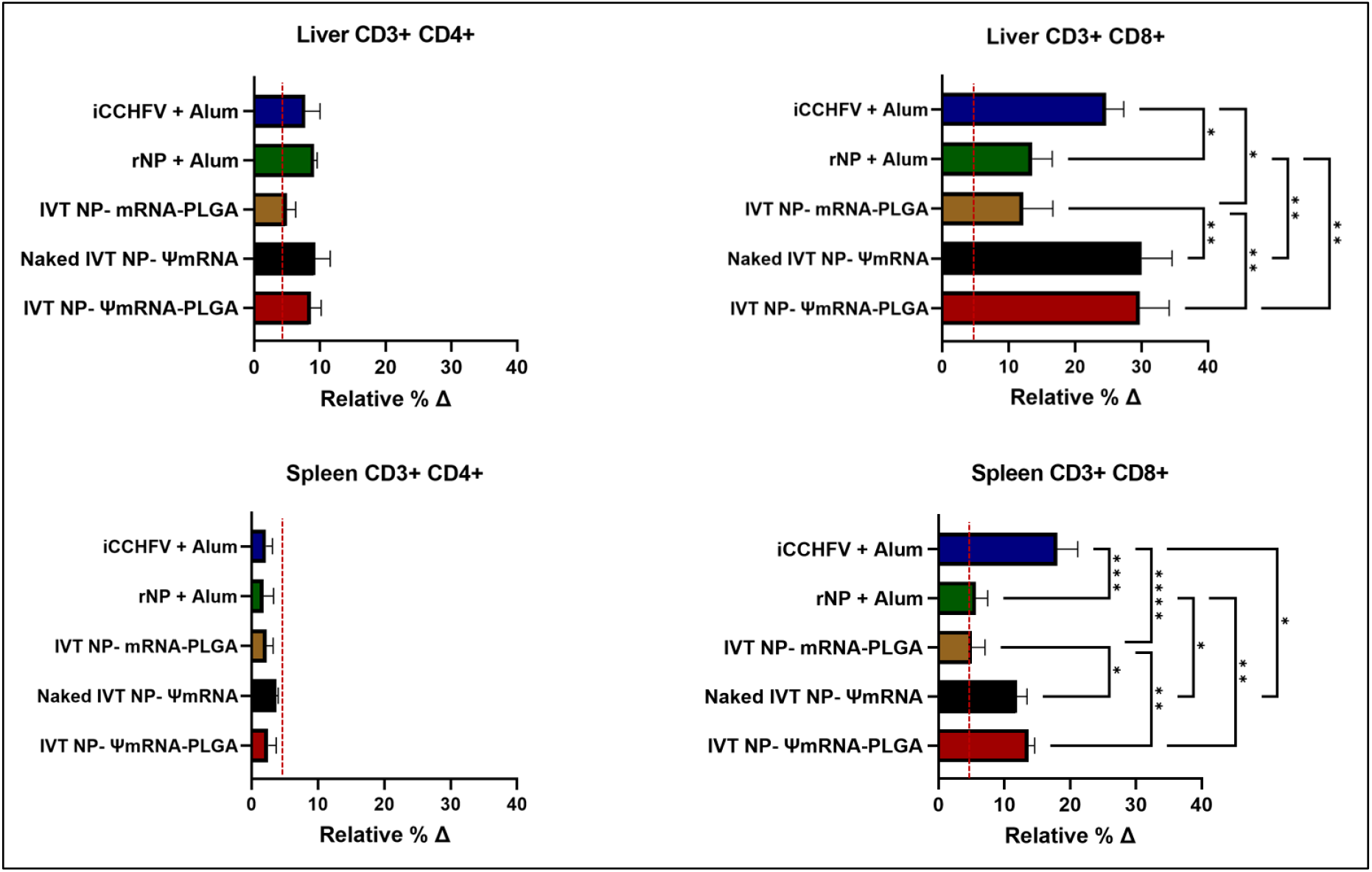
Enhanced CD8+ T Cell Profile in Surviving Mice Post-Challenge. Flow cytometry analysis of spleen and liver tissues showed a notable increase in CD8+ T cells, up to 30%, especially in mouse groups with a 100% survival rate at 2 dpi.

### Analysis of Cellular Immunity: Cytokine Response

To assess the elements of the cellular immunity, the gene expression levels of specific cytokines were determined in spleen and liver cells using qPCR analyses (+2 DPI). As shown in Figure 9, significant increases in IFN-γ, TNF-α, IL-10, IL-12, and IL-2 levels were detected in the liver and spleen compared to the Sham group. These increases were more pronounced in mouse groups with high survival rates. It was observed that IL-15, GM-CSF, and IL-4 levels rose particularly in the liver, while IL-10, IL-6, and IL-5 levels increased in IVT NP-ΨmRNAs groups. IL-1β was significantly elevated only in the liver of mice immunized with iCCHFV, whereas IL-17A was observed solely in the spleen. After cell sorting, the cytokine responses in CD3+CD8+ and CD3+CD4+ cells in the spleen and liver were also examined, revealing that IFN-γ, TNF-α, and IL-4 were closely associated with survival rates (Figure S11) ((appendix pp 19-20).

**Figure 9.**
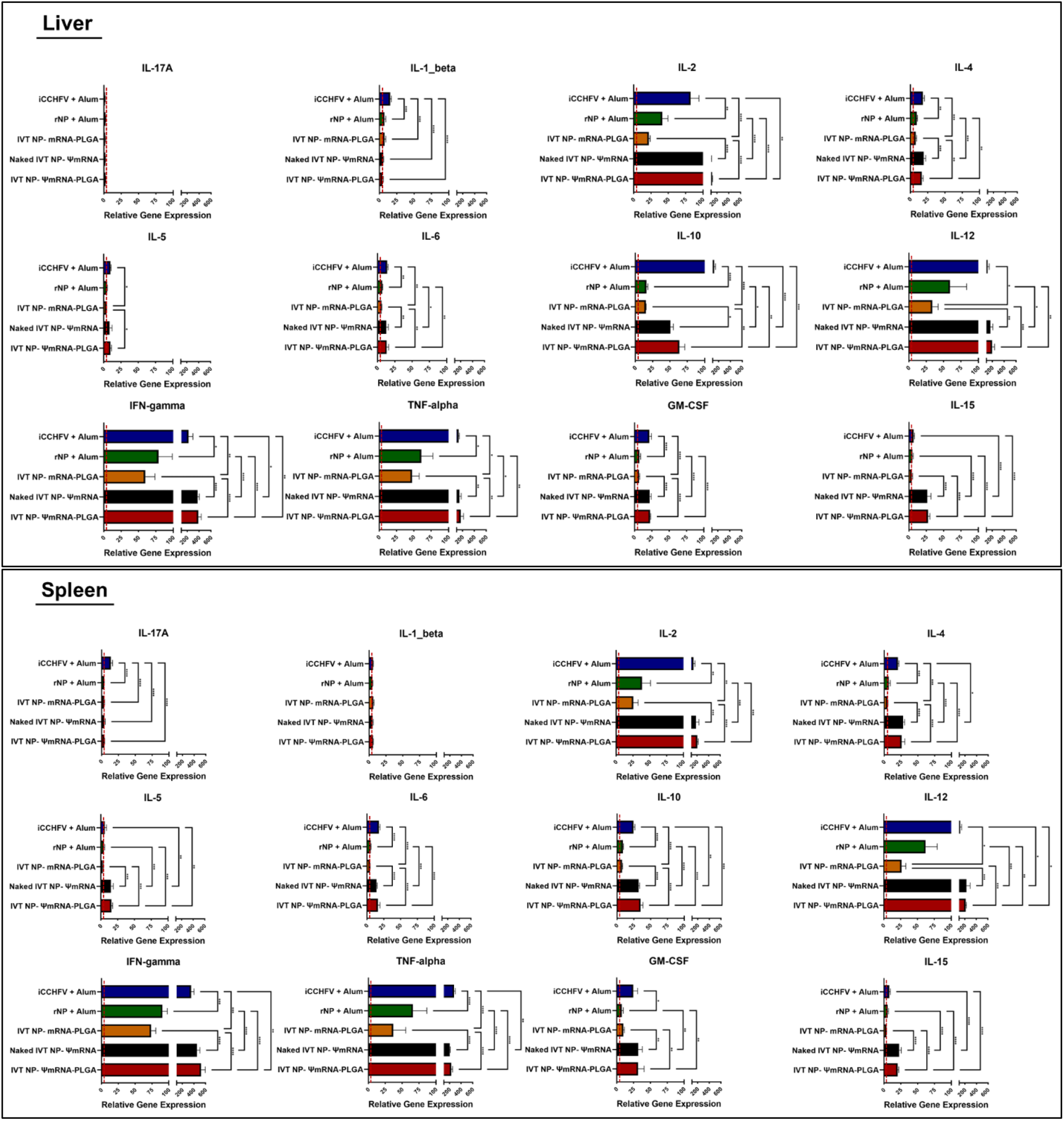
Cytokine Profiles in Liver and Spleen Cells Post-Immunization. qPCR analysis demonstrated significant increases in IFN-γ, TNF-α, IL-12, and IL-2 levels in liver and spleen tissues of immunized mice compared to the Sham group, with more pronounced increases in groups exhibiting high survival rates. Elevated levels of IL-15, GM-CSF, and IL-4 were particularly noted in the liver, while IL-10, IL-6, and IL-5 levels rose in IVT NP-ΨmRNA groups. IL-1β was significantly elevated only in the liver of iCCHFV-immunized mice, and IL-17A was detected solely in the spleen.

## Discussion

The most remarkable finding of this study is the full protection observed in mice against a lethal challenge of CCHFV following immunization with IVT NP-ΨmRNAs encoding NP only.

The report represents a part of a broader scrutiny into the potential of modified mRNA-based vaccines to elicit immune responses against various viral pathogens and tumors. The success of mRNA vaccines developed for COVID-19 has generated significant interest in applying this technology to other pathogens. Achieving a robust immune response against lethal viruses like CCHFV is critical for public health in a large part of the world(26, 32–36). Previously, several vaccine candidates have been explored, including inactivated virus formulations, vector-based vaccines, and even mRNA vaccines encoding CCHFV proteins(13, 18, 30, 34, 37–39). The results of this study demonstrated that NP alone, when presented as a modified mRNA vaccine, could afford complete protection against lethal challenge, therefore, pointing to the potential of a vaccine candidate comprised of a single antigen for CCHFV infection (Figure 5). This might represent a welcome data for regions affected by CCHF for which no approved vaccine formulation and antiviral treatment with proven antiviral efficacy exist.

Our study provides compelling evidence that IVT-mRNA vaccination encoding the CCHFV NP antigen elicits a robust immune response, demonstrating both humoral and cellular immunity against CCHFV infection. While previous studies have explored alternative RNA-based platforms, such as alphavirus-derived replicating RNA (repRNA) vaccines(40, 41), our findings highlight the effectiveness of a non-replicating IVT-mRNA approach. Compared to previous reports in which a single 100 ng dose of repRNA encoding NP or a combination of NP and GPC was used, our study employed IVT-mRNA encoding NP alone, which was sufficient to trigger a completely protective immune response with a significant induction of IFN-γ-producing T cells.

In other reports, non-human primates (NHPs) were utilized as model organisms, where DNA vaccines and repRNA platforms using NP as the primary antigen demonstrated that NP-based immunization elicits strong but non-neutralizing antibody responses, contributing to protection(36, 42). These studies also indicated that GPC-specific T cells play a pivotal role in viral clearance. Interestingly, our results align with this concept, as we observed a predominant Th1-skewed immune response, further supporting the notion that cellular immunity is a key determinant of protection against CCHFV. Moreover, it has been suggested that IFN-γ serves as a crucial antiviral cytokine during acute CCHFV infection(43). Our data reinforce this hypothesis, as a marked increase in IFN-γ expression was detected post-immunization, emphasizing the protective role of T-cell responses in mRNA vaccine-induced immunity.

The question of whether an internal viral protein can afford protective humoral antiviral immunity has recently been addressed(44). Accordingly, a TRIM21-mediated mechanism plays a key role in antibody-mediated protection, where the intracellular Fc receptor TRIM21 is functional. Whether a similar mechanistic role is played by TRIM21 in our model remains to be addressed.

The in vitro transcripts of Ψ-modified mRNA are reported to have increased stability and led to minimized innate immune responses(45, 46). In our study, we have shown that IVT Ψ-modified mRNA expressed abundantly in transfected Huh-7 cells, as demonstrated in Western blot, EIA, and immunofluorescence assays (Figure 2). The temporal analysis showed that up to the 48th hour post-transfection, the cells continued to synthesize the viral protein from transfected templates, pointing to the stability of the mRNAs. More significantly, however, Ψ-modified mRNA induced fully protective immunity against lethal viral challenge compared to unmodified mRNA (Figure 5), therefore confirming the previously published extensive data on the increased immunogenicity of engineered mRNA containing Ψ(11, 13, 47).

After the production of IVT mRNA, the vaccines were formulated with PLGA nanoparticles to enhance mRNA stability and resistance to enzymatic degradation. PLGA is also reported to improve immune responses by enhancing controlled release and biodistribution(16, 27, 29). In this study, PLGA-nanoparticle-formulated vaccine candidate (IVT NP-ΨmRNA-PLGA) did not induce significantly stronger immune responses compared to naked IVT NP-ΨmRNA. The full impact of PLGA on NP vaccinations could be assessed in a setting where different concentrations of antigens are utilized.

Inactive viral antigens iCCHFV introduced in aluminum hydroxide, a commonly used human adjuvant, were also effective against viral challenge (Figure 5), confirming previously reported data where full virions could protect against viral challenges(18). When NP alone is introduced to the immune system as a protein antigen, it appears that the protection is diminished significantly to 40% (Figure 5). These findings are not surprising given the many previous reports on the incomplete immunity acquired following NP immunizations as a single antigen(13, 23, 39, 48, 49).

In this study, we also evaluated both humoral and cellular immune responses in the immunized animals. Humoral immune responses, evaluated using serum EIA tests, showed high NP-specific IgG antibody titers in mice immunized with IVT NP-ΨmRNAs (Figure 3 and 7). These results demonstrate that the vaccine generated a robust humoral response in both BALB/c and C57BL/6 mouse models. Lymphoproliferation and flow cytometry analyses confirmed that the vaccines also induced a strong cellular immune response against CCHFV, with an increased CD8+ T-cell population in the liver and spleen, correlating with the vaccine’s protective efficacy.

Challenge experiments revealed that both the IVT NP-ΨmRNA-PLGA formulation and naked IVT NP-ΨmRNA provided 100% survival rates and significantly reduced viral loads in the liver and spleen (Figure 6). These findings suggest that Ψ-modified mRNA vaccines could reduce the number of viruses in visceral organs, which might have critical roles in the development of fatal hemorrhagic fever. Comparing the results from the PLGA-encapsulated IVT NP-ΨmRNA group with naked ΨmRNA test subjects, we noticed that the difference was not striking. It is likely that additional studies with different encapsulation methods, such as those with other lipid nanoparticles, might provide further information on this question.

Cytokine response analysis showed that IVT NP-ΨmRNAs immunizations increased the production of pro-inflammatory cytokines such as IFN-γ, TNF-α, IL-2, and IL-12. These cytokines are also involved in promoting cellular immune responses. The elevated levels of these cytokines could indicate that IVT NP-ΨmRNAs immunization might favor a strong cellular and inflammatory response against viral infection. Obviously, these points need to be scrutinized in more detail.

In conclusion, our IVT NP-ΨmRNA vaccine candidate demonstrated complete protection against CCHFV by inducing both humoral and cellular immune responses, with PLGA-nanoparticle formulations highlighting their potential as an effective strategy for combating future viral threats. These findings, in conjunction with previous studies, underscore the importance of NP as a promising target for CCHFV vaccine development and reaffirm the potential of Ψ-modified mRNA vaccines to generate strong immune responses against viral hemorrhagic fever agents. Moreover, our IVT-mRNA platform offers a safer and more scalable alternative to existing DNA and replicating RNA-based approaches. Future studies should focus on assessing the durability and breadth of immune responses, as well as conducting challenge experiments in higher animal models, to facilitate the clinical translation of NP-based mRNA vaccines.

